# Bacterial CRISPR-Cas Abundance Increases Precipitously at Around 45°C: Linking Antivirus Immunity to Grazing Risk

**DOI:** 10.1101/2021.05.25.445389

**Authors:** Xin-Ran Lan, Zhi-Ling Liu, Deng-Ke Niu

## Abstract

Although performing adaptive immunity, CRISPR-Cas systems are present in only 40% of bacterial genomes. Here, we observed an abrupt transition of bacterial CRISPR-Cas abundance at around 45°C. Phylogenetic comparative analyses confirmed that the abundance correlates with growth temperature only at the temperature range around 45°C. Meanwhile, we noticed that the diversities of cellular predators have a precipitous decline at this temperature range. The grazing risk faced by bacteria reduces substantially at around 45°C and almost disappears above 60°C. So viral lysis would become the dominating factor of bacterial mortality, and antivirus immunity has a higher priority. In temperature ranges where the abundance of cellular predators does not change with temperature, temperatures would have negligible effects on CRISPR-Cas abundance. The hypothesis predicts that bacteria should also be rich in CRISPR-Cas systems if they live in other extreme conditions inaccessible to grazing predators.

## Introduction

CRISPR-Cas is the adaptive immune system of bacteria and archaea. Like the adaptive immune system of jawed vertebrates, it can remember previously encountered pathogens, initiate a rapid response to a second invasion, and eliminate the recurrent invader. However, unlike the ubiquitous presence of adaptive immunity in jawed vertebrates,^1^ the CRISPR-Cas systems are only present in about 40% of bacteria.^2^ The patchy distribution of the CRISPR-Cas systems among bacteria is a recognized mystery.^3, 4^ Given the constant horizontal transfers, their absence in more than half of bacterial genomes is unlikely to happen just by chance.^5^ Instead, it should be attributed to a tradeoff between the costs and benefits of the CRISPR-Cas systems. First, the acquirement and maintenance of CRISPR-Cas systems would sequestrate limiting resources such as the building blocks, the energy, and the transcription and translation machines.^6-8^ Second, the autoimmune response and cell death induced by self- and prophage-targeting spacers might be a selective force for the loss of CRISPR-Cas systems.^9, 10^ In addition, for the viruses with high densities, high mutation rates, high genetic diversities, or carrying anti-CRISPR proteins, the efficiency of CRISPR-Cas systems is limited.^11-14^ CRISPR-Cas systems are not favored in conditions with high antibiotic pressures because they inhibit horizontal gene transfer, an efficient way for bacteria to acquire antibiotic resistance.^15^

Just after discovering the CRISPR-Cas structures, it was noticed that they are more prevalent in the thermophilic archaea and the hyperthermophilic bacteria.^16, 17^ Later large-scale analyses confirmed the prevalence of CRISPR-Cas systems in thermophiles and hyperthermophiles and showed a positive correlation between CRISPR abundance and a growth temperature.^13, 18-20^ Recently, a computer learning approach with an attempt to control the phylogenetic bias suggests that temperature and oxygen levels are the most influencing ecological factors determining the distribution of the CRISPR-Cas systems.^21^ In this study, we carried out a detailed analysis on the thermal distribution of CRISPR-Cas systems and observed a precipitous increase of bacterial CRISPR-Cas systems at around 45°C. At temperatures higher and lower than this range, bacterial CRISPR-Cas abundance does not correlate with growth temperature. The association of adaptive immunity with a narrow range of growth temperature provokes a new hypothesis linking the antivirus immunity of bacteria to their grazing risk by predatory protists and predatory bacteria.

## Materials and Methods

From the growth temperature database TEMPURA and the Genome Taxonomy Database,^22, 23^ we retrieved 682 bacteria and 156 archaea with growth temperatures and phylogenetic information. The sequences of these genomes were downloaded from ftp://ftp.ncbi.nlm.nih.gov/genomes/. We annotated their CRISPR-Cas systems using CRISPRCasFinder v1.3 program.^24^ The raw data of these 682 bacteria and 156 archaea were deposited as Table S1.

According to Couvin et al.,^24^ the annotated CRISPR arrays were classified into four categories, 1 to 4, according to their evidence levels. The CRISPR arrays with evidence levels 3 and 4 are highly likely candidates, and those with evidence levels 1 and 2 are potentially invalid. Therefore, only the CRISPR arrays with evidence levels 3 and 4 were counted in calculating CRISPR array abundance. We counted the putative CRISPR arrays with evidence levels 1 and 2 as zero and presented the results in the main text and Table S2. We also replicated our analyses by regarding the putative CRISPR arrays with evidence levels 1 and 2 as controversial CRISPR arrays and discarding the species having only CRISPR arrays with evidence levels 1 and 2 in calculating CRISPR array abundance. That is, these species were excluded from both numerator and denominator in the calculation of CRISPR-Cas abundance. Similar results (Table S3) were obtained, and the same conclusion was supported.

The CRISPR array abundance in a bacterial or archaeal group was defined as the total number of CRISPR arrays annotated in their genomes divided by the genome numbers of the group. The abundances of CRISPR spacers, *cas* genes, and *cas* gene clusters were defined similarly.

The phylogenetic signals (λ) of CRISPR array abundance, CRISPR spacer abundance, *cas* gene abundance, *cas* gene cluster abundance, and growth temperatures were estimated using the phylosig function of the R (Version 4.0.3) package phytools (Version 0.7-70)^25^. The phylogenetic generalized least squares (PGLS) regression was performed using the R (Version 4.0.3) package phylolm (version 2.6.2)^26^. Pagel’s lambda model has been applied in the analyses.

## Results

### Bacterial CRISPR array abundance jumps up at around 45°C

By plotting the CRISPR array abundance against the optimal growth temperatures (Topt) in a column chart, we see a novel pattern on the thermal distribution of CRISPRs in bacteria (Fig. 1A). An abrupt transition of bacterial CRISPR array abundance happens at around 45°C. The CRISPR array abundance also fluctuates below and above 45°C, but the amplitudes of the fluctuations are much weaker. By sliding the windows of each column and calculating the ratio of CRISPR array abundance of one column to the next, we confirmed that the transition of CRISPR array abundance at 45°C is the most sudden one (Fig. 1B).

**Fig. 1.**
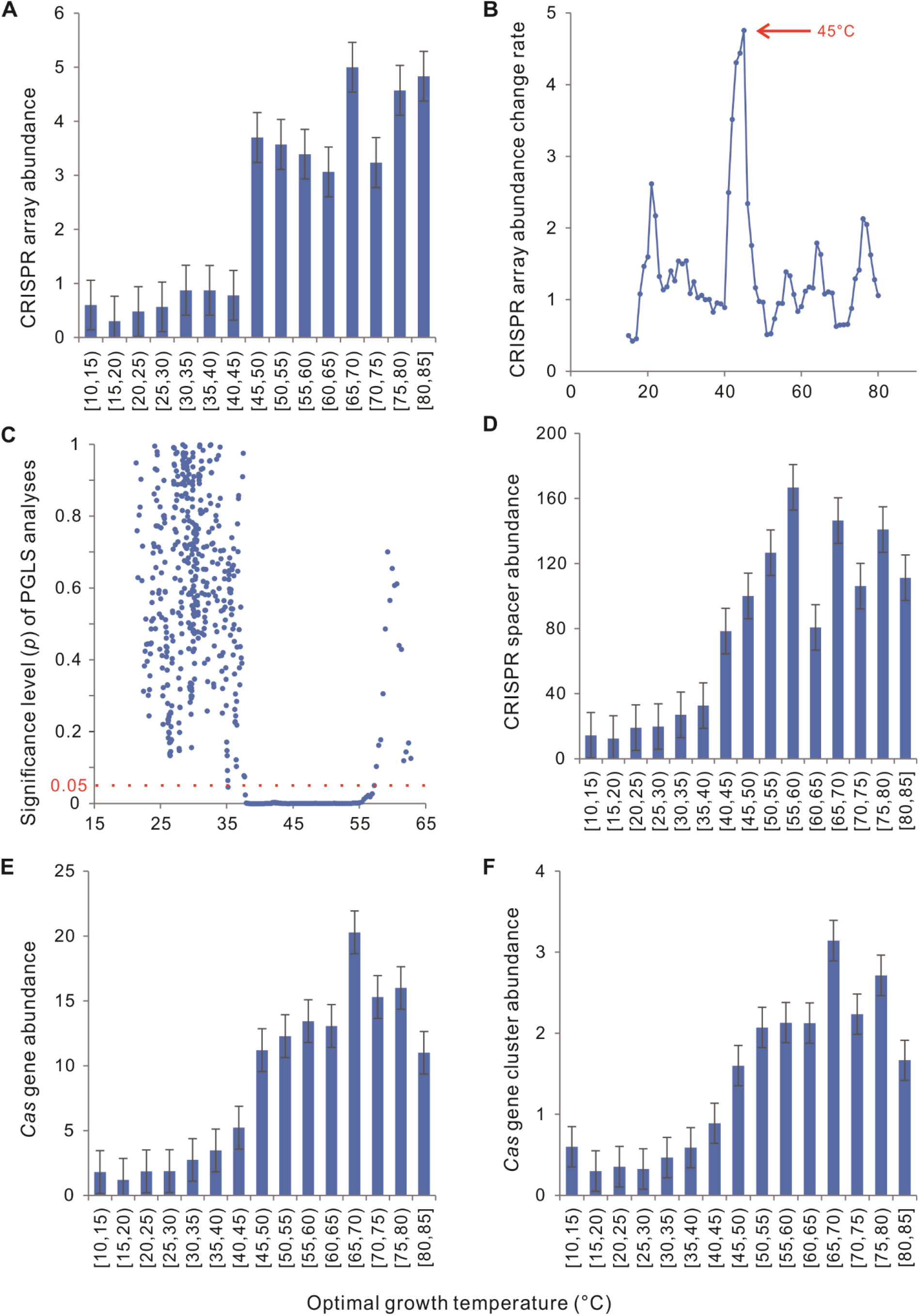
Distinctive relationships of bacterial CRISPR-Cas abundance with different optimal growth temperature (Topt) ranges. (A) An abrupt jump up of bacterial CRISPR array abundance occurs at around 45°C. (B) The change rate of bacterial CRISPR array abundances along the axis of Topt, which was calculated from the ratio of bacterial CRISPR array abundance of each column to the next. The columns not shown in Fig. 1A were obtained by sliding one degree for each time. (C) Phylogenetic generalized least squares (PGLS) analysis showed significant positive correlations between bacterial CRISPR array abundances and Topt only around 45°C. The 682 bacteria were aligned along the Topt axis. One hundred neighboring samples were taken in each round of PGLS analysis. (D-F) The abundances of CRISPR spacer, *cas* genes, and *cas* gene clusters increase substantially with Topt at around 45°C, but no abrupt jumps have been observed. In (A) and (D-F), the average values and the standard errors are presented.

For a quantitative measure for the relationship between bacterial CRISPR array abundance and Topt, we first measured their phylogenetic signals (λ = 0.85, *p* = 3 × 10^−44^ for CRISPR array abundance and λ = 0.95, *p* = 4 × 10^−197^ for Topt) and proved the necessity of using phylogenetic comparative methods to control the effects of common ancestors ^27, 28^. Their correlation was examined by the phylogenetic generalized least squares (PGLS) regression, where a significant positive slope corresponds to a significant positive correlation, and a negative slope indicates the reverse. Globally, bacterial CRISPR array abundance is positively correlated with Topt (slope = 0.06, *p* = 8 × 10^−12^). However, when the bacteria were classified into three categories according to their Topts, low temperatures (10 ≤ Topt ≤ 35°C, *n* = 488), moderate temperatures (35 < Topt < 50°C, *n* = 97), and high temperatures (50 ≤ Topt ≤ 85°C, *n* = 97), a significant correlation was only observed in the moderate-temperature bacteria (Table 1). Furthermore, we aligned the 682 bacteria along the Topt axis and performed a PGLS analysis for every 100 neighboring samples. Significant correlations were observed only in a range around 45°C (Fig. 1C).

**Table 1.** Correlations between bacterial CRISPR-Cas abundance and optimal growth temperature

|  |  | CRISPR<br>array | CRISPR<br>spacer | *Cas*<br>gene | *Cas* gene<br>cluster |
| --- | --- | --- | --- | --- | --- |
| Bacteria |  |  |  |  |  |
| 10–35°C (*N*=488) | Slope | 0.020 | −0.102 | 0.030 | 0.001 |
|  | *p* | 0.297 | 0.771 | 0.475 | 0.866 |
| 36–48°C (*N*=97) | Slope | 0.239 | 6.338 | 0.855 | 0.118 |
|  | *p* | **2×10<sup>−4</sup>** | **3×10<sup>−4</sup>** | **10<sup>−5</sup>** | **10<sup>−4</sup>** |
| 50–85°C (*N*=97) | Slope | 0.045 | −0.302 | 0.061 | 0.009 |
|  | *p* | 0.171 | 0.851 | 0.613 | 0.660 |
| 10–85°C (*N*=682) | Slope | 0.065 | 1.598 | 0.035 | 0.220 |
|  | *p* | **8×10<sup>−12</sup>** | **2×10<sup>−6</sup>** | **4×10<sup>−14</sup>** | **3×10<sup>−15</sup>** |
| Archaea |  |  |  |  |  |
| 18–41°C (*N*=52) | Slope | −0.015 | 0.613 | 0.021 | 0.136 |
|  | *p* | 0.752 | 0.783 | 0.416 | 0.469 |
| 41.5–82.5°C (*N*=53) | Slope | 0.117 | 1.639 | 0.003 | 0.007 |
|  | *p* | **0.008** | 0.194 | 0.832 | 0.943 |
| 83–106°C (*N*=51) | Slope | 0.141 | 2.250 | 0.372 | 0.075 |
|  | *p* | 0.139 | 0.214 | **0.046** | **0.009** |
| 18–106°C (*N*=156) | Slope | 0.097 | 0.912 | 0.021 | 0.107 |
|  | *p* | **2×10<sup>−6</sup>** | 0.100 | **0.002** | **0.012** |

### Bacterial CRISPR spacer abundance increases precipitously at around 45°C

Globally, bacterial CRISPR spacer abundance correlates positively with Topt (*p* = 2 × 10^−6^, Table 1). Although there is no abrupt jump in bacterial CRISPR spacer abundance along the axis of growth temperature, the spacer abundance increases substantially with Topt only at around 45°C (Fig. 1D). PGLS analyses showed a significant correlation between CRISPR spacer abundance and Topt at the moderate temperatures, but not in the low temperatures or the high temperatures (Table 1).

### Bacterial *cas* gene abundance increases precipitously at around 45°C

Across all the 682 bacteria with Topt ranging from 10−85°C, both *cas* gene abundance and *cas* gene cluster abundance are significantly correlated with Topt (Table 1). An abrupt jump at around 45°C could be observed in bacterial *cas* gene cluster abundance, and the increase in bacterial *cas* gene and *cas* gene cluster abundances are precipitous at around 45°C (Fig. 1E-F). PGLS analyses gave the same statistical conclusion for *cas* gene abundance and *cas* gene cluster abundance. Both abundances are significantly correlated with Topt at the moderate temperatures (*p* = 3 × 10^−4^), but not in the low temperatures (*p* = 0.77) or the high temperatures (*p* = 0.85) (Table 1).

### Archaeal CRISPR-Cas abundance has a less distinctive pattern

We also examined the relationship between growth temperature and CRISPR-Cas abundance in archaea. Globally, archaeal Topt positively correlates with CRISPR array abundance, *cas* gene abundance, and *cas* gene cluster abundance, but not with spacer abundance (Table 1). However, no abrupt jumps of CRISPR-Cas abundance were observed in archaea (Fig. 2). It should be noted that the sample sizes of some columns are tiny, and so the observed pattern is sensitive to the presence of a few outliers. For example, in the temperature range [25,30), there are only two archaeal species, *Methanosarcina lacustris* and *Methanosphaerula palustris*, with eight and one CRISPR arrays, respectively.

**Fig. 2.**
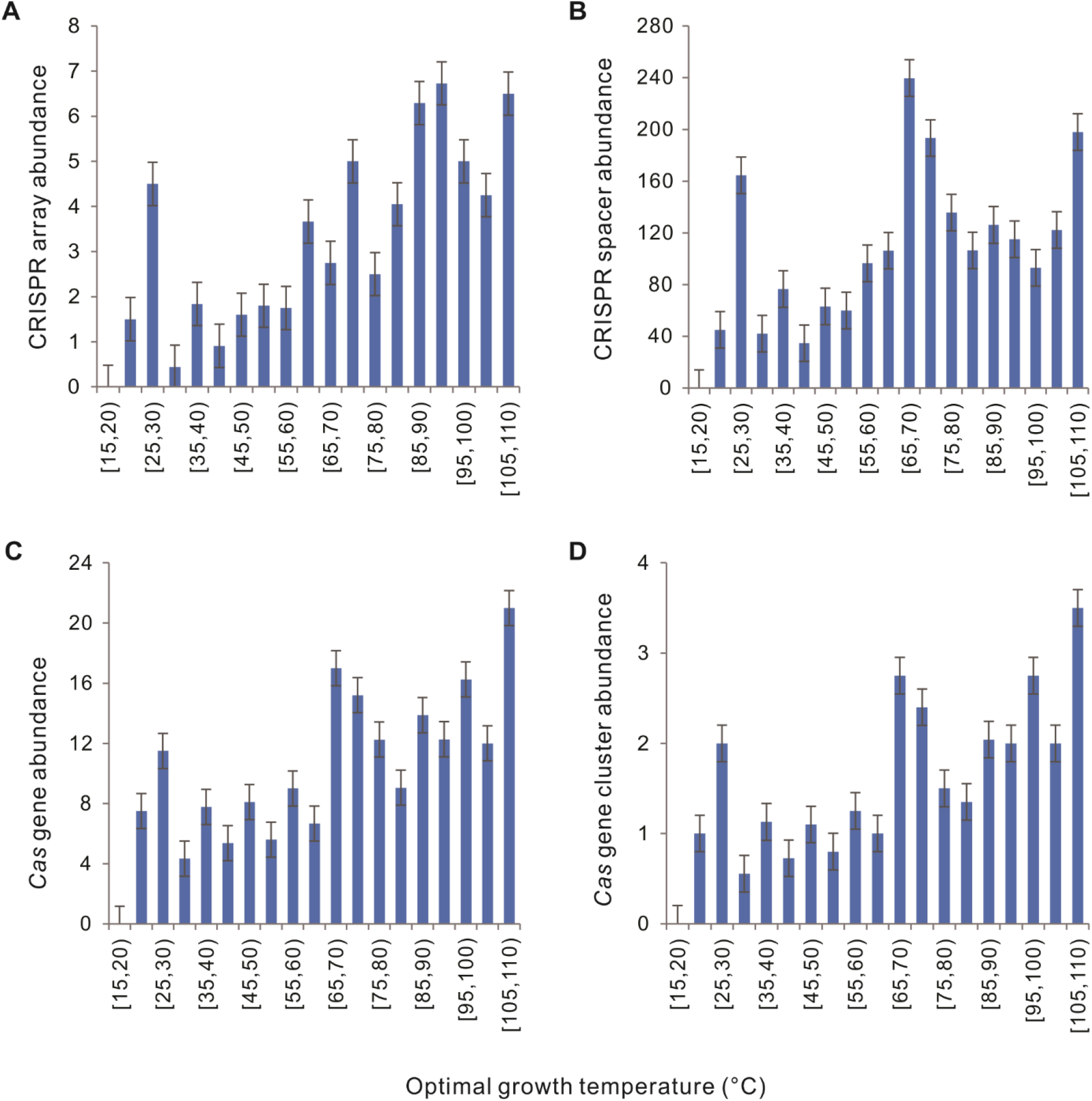
Relationships of archaeal CRISPR-Cas abundance with different optimal growth temperature (Topt) ranges. (A) CRISPR arrays. (B) CRISPR spacers. (C) *cas* genes. (D) *cas* gene clusters. The average values and the standard errors are presented.

Because the 156 archaeal species are enriched in thermophiles and hyperthermophiles, dividing them into three temperature range categories using the above thresholds leads to a too-small sample for the low-temperature category (*N* = 17). Therefore, we divided the 156 archaea into three nearly equal-sized groups according to their Topts (Table 1). A significant correlation between Topt and CRISPR array abundance could be observed in the moderate-temperature group (*n* = 53, 41.5 ≤ Topt ≤ 82.5°C, slope = 0.12, *p* = 0.01), but not in the lower (*p* = 0.75) or the higher one (*p* = 0.14). The CRISPR spacer abundance is not correlated with Topt in all three groups. The *cas* gene abundance and *cas* gene cluster abundance are positively correlated with Topt only in the high-temperature group (Table 1).

### No significant difference in CRISPR-Cas abundance between bacteria and archaea living at the same temperature

We rounded up the values of Topts to integers and selected the temperatures that are Topts of both bacterial species and archaeal species. The CRISPR-Cas abundance of bacteria/archaea with the same Topt were averaged. Pairwise comparison of the obtained 35 archaeal-bacterial pairs did not show any significant difference in CRISPR array abundance, spacer abundance, *cas* gene abundance, or *cas* gene cluster abundance (Wilcoxon signed ranks test, *p* = 0.588, 0.406, 0.447, and 0.925, respectively).

## Discussion

Previous studies have found that prokaryotes living in high temperatures are more likely to have CRISPR-Cas systems.^13, 16-21^ In this study, we presented a more detailed description of the relationship between CRISPR-Cas and Topt. In low and high temperatures, the abundances of bacterial CRISPR-Cas are not significantly correlated with Topt. However, at around 45°C, bacterial CRISPR-Cas abundance increases sharply with the increase of Topt. Most significantly, the bacterial CRISPR array abundance exhibits an abrupt jump at 45°C. As we see, the evolutionary and mechanical links between growth temperature and CRISPR-Cas abundance previously proposed^13, 14, 29^ could not explain the abrupt transition at around 45°C. Temperature influences many aspects of cellular processes and the physical features of the environment.^30^ No matter linear or nonlinear, the physical effects of temperature increase gradually with temperature.

We noticed that the thermal distribution of eukaryotes has an abrupt decline at around 45°C. Very few members of each group of eukaryotes can live above 60°C ^30-33^. Some bacteria could also kill other bacteria and consume the released nutrients. Except for a few exceptions, most predatory bacteria could not grow at temperatures above 45°C.^34, 35^ Here, we propose that the accessibility of cellular predators along the temperature axis might indirectly govern the thermal distribution of the CRISPR-Cas systems (Fig. 3).

**Fig. 3.**
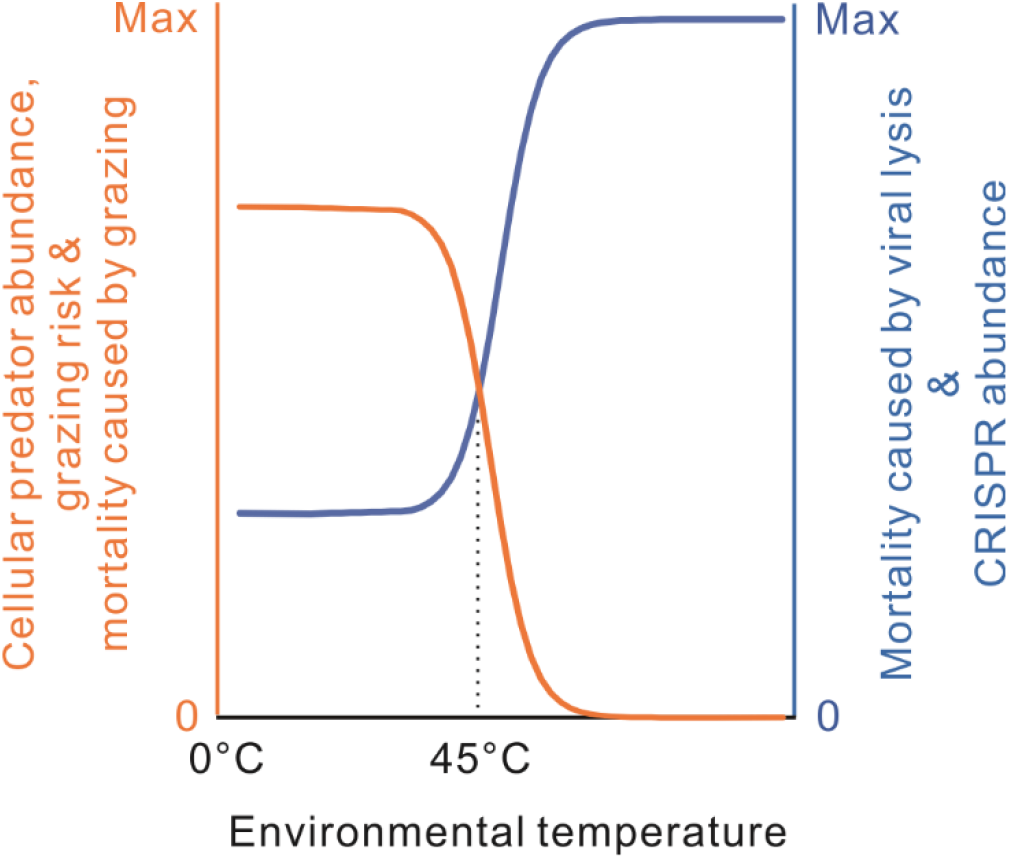
A model for the thermal distribution of bacterial CRISPR-Cas systems. At low temperatures, the evolution of CRISPR-Cas systems is limited by the grazing pressure of cellular predators. At around 45°C, cellular predator abundance decreases abruptly with environmental temperature, so viral lysis takes more in bacterial mortality, and the requirement of the immune function increases substantially. When cellular predators almost disappear at high temperatures, mortality results primarily from viral lysis; most bacteria evolve the highest antivirus capacity.

In birds, it has been shown that predation risk could significantly reduce the allocation of the limiting resources to immune function.^36^ The birds captured by cats consistently had smaller spleens than those killed by nonpredatory reasons like collisions with windows or cars. When the hosts are exposed to lethal predators, predator-mediated mortality becomes dominant, and the pathogen-mediated mortality decreases relatively. Consequently, the priority to invest in immune function is reduced. Physiological and evolutionary reducing the allocation of the limiting resources to immune function would be favored.

We proposed that the same case might happen in bacteria. Most of the bacterial mortality is caused by cellular predator grazing and viral lysis.^37^ For picophytoplankton, grazer-mediated mortality and viral-mediated mortality are inversely correlated.^38, 39^ Indirect interactions among grazers and viruses are destined to occur provided that bacterial cells have tradeoffs in the grazing resistance and virus resistance. When bacterial mortality mostly comes from predator grazing, the benefits of adaptive immunity might be dwarfed by the costs in the maintenance and expression of the CRISPR-Cas systems, like allocating the limiting resources, targeting host or prophage genome, and inhibiting horizontal gene transfer. By contrast, when bacterial mortality mostly comes from viral lysis, some costs of CRISPR-Cas systems would be tolerable because the benefits of adaptive immunity outweigh the costs. At around 45°C, with the increase of environmental temperature, cellular predator abundance decreases sharply;^30, 32^ thus, the grazing risk of bacteria is abruptly relieved. Bacteria growing at higher temperatures mainly die from viral lysis. Antivirus immunity is favored even if it has some costs on the bacterial cells. Within the temperature ranges inaccessible to grazing predators or within the temperature ranges fully accessible to the grazing predators, temperature changes have little effect on the evolution of the antivirus system (Fig. 3).

We propose that bacterial cells tend to lose the CRISPR-Cas systems when bacterial mortality mostly comes from predator grazing. In natural environments, however, this does not always happen even below 45°C. Bacterial mortality is destined to be more or less contributed by viral lysis because of the widespread of viruses. In addition, prey bacteria are not entirely passive to be grazed, and they could evolve grazing-resistance capacities, like high motility, large size, and biofilm formation.^40-42^ In a long-term arms race between prey bacteria and grazing protists/bacteria, the prey bacteria might, in some periods, be free of grazing risk and grazing-caused mortality because of newly evolved grazing-resistance strategies. In this case, viral lysis becomes dominant in bacterial mortality, and the bacteria experience an intense selective pressure to have CRISPR-Cas systems. Therefore, our hypothesis is not exclusive to CRISPR-Cas-rich psychrophilic and mesophilic bacteria.

In addition, the new hypothesis is to explain the abrupt transition of CRISPR-Cas abundance at around 45°C. Besides growth temperature, many other ecological and evolutionary factors that might influence the phylogenetic distribution of CRISPR-Cas systems^8-15^ are beyond the scope of this hypothesis. Even for the thermal distribution of CRISPR-Cas systems, we are open to other possible explanations. A bacteriophage infecting the tropical pathogen *Burkholderia pseudomallei* was found to be temperate at lower temperatures (25°C) and tends to go through a lytic cycle at higher temperatures (37°C).^43^ At least for type I CRISPR-Cas systems, targeting temperate phages has been demonstrated to be a driving force for the loss of adaptive immunity.^10^ Therefore, bacteria living in low temperatures might have fewer CRISPR-Cas systems because of the temperate-phage-induced bacterial autoimmunity. However, the temperature-associated switching of the life cycle of the bacteriophage of *B. pseudomallei* is just a piece of isolated evidence. At present, we do not know how many phages in nature have similar temperature-associated switching of life cycle as the bacteriophage of *B. pseudomallei*.

Besides explaining the thermal distribution, our hypothesis suggests that other environmental factors that severely reduce cellular predator abundance should also affect bacterial mortality and CRISPR-Cas distribution. A generalized prediction of our hypothesis is that bacteria living in extreme environments inaccessible to cellular grazers should carry more CRISPR-Cas systems in their genomes.

Weissman et al. recently found a negative interaction between CRISPR-Cas systems and oxygen availability and hypothesize that oxidative-stress-associated DNA repair processes might interfere with the function of CRISPR-Cas systems.^21^ Here, we provide an alternate explanation for their observation by extending our hypothesis. All the well-known cellular predators are aerobic organisms. In anoxic environments, there are no cellular predators or only a few unknown predators. Similar to growth temperatures at around 45°C, the transition from an oxic environment to an anoxic environment would substantially reduce the grazing-caused bacterial mortality and indirectly increase the requirement of antivirus immunity.

There is no abrupt jump in CRISPR-Cas abundance in archaea along the axis of growth temperature (Fig. 2). We found that Topt and CRISPR array abundance are positively correlated in the moderate temperatures (41.5 ≤ Topt ≤ 82.5°C), but not in the lower or higher temperatures (Table 1). However, similar patterns have not been observed in the CRISPR spacer, *cas* gene, or *cas* gene cluster (Table 1). The difference in CRISPR-Cas distribution between bacteria and archaea might come from the physiological, genomic, or ecological differences between the two domains. It is also possible that random noises resulting from the small sample size have masked the thermal distribution pattern of CRISPR-Cas among archaea. In a recent analysis on the relationship between growth temperature and GC content using the same growth temperature dataset,^22^ we found that Topt is significantly correlated with bacterial genome GC content (*N* = 681) but not archaeal genome GC content (*N* = 155). Then, we randomly drew 155 bacteria from the 681 bacteria 1000 times. In > 95% rounds of resamplings, the positive correlations became statistically nonsignificant (*P* > 0.05).^44^ The results suggest that the effective sample size in phylogenetically-related data is much smaller than the census number of the analyzed lineages. We hope to replicate the present analysis in archaeal genomes in the future when several hundred or thousands of archaeal species are available for analysis.

## Conclusion

The CRISPR-Cas systems are known to be enriched in thermophilic and hyperthermophilic prokaryotes. In this paper, we take a step further by revealing an abrupt transition of bacterial CRISPR-Cas abundance at around 45°C and putting forward a new hypothesis on the thermal distribution of bacterial CRISPR-Cas systems. Grazing of cellular predators and viral lysis are the primary sources of bacterial mortality; their negative interaction largely influences the tradeoffs between the costs and benefits of antivirus strategies and grazing resistance strategies. As cellular predator diversities and grazing risk precipitously decline at around 45°C, viral lysis becomes the dominant source of bacteria mortailty, and the requirement of adaptive immunity is increased indirectly.

## Acknowledgments

We thank Quan-Guo Zhang and Wen-Hong Deng for helpful discussions and Christine Pourcel and Pierre-Albert Charbit for technical supports.

## Authorship confirmation statement

DKN conceived the study and wrote the manuscript. XRL and ZLL performed the analysis. All co-authors have reviewed and approved of the manuscript prior to submission.

